# Interconversion between Anticipatory and Active GID E3 Ubiquitin Ligase Conformations via Metabolically Driven Substrate Receptor Assembly

**DOI:** 10.1101/824060

**Authors:** Shuai Qiao, Christine R. Langlois, Jakub Chrustowicz, Dawafuti Sherpa, Ozge Karayel, Fynn M. Hansen, Viola Beier, Susanne von Gronau, Daniel Bollschweiler, Tillman Schäfer, Arno F. Alpi, Matthias Mann, J. Rajan Prabu, Brenda A. Schulman

**Author notes:** Equal contributions (Christine R. Langlois, Jakub Chrustowicz, Dawafuti Sherpa).

## Abstract

Cells respond to environmental changes by toggling metabolic pathways, preparing for homeostasis, and anticipating future stresses. For example, in *Saccharomyces cerevisiae*, carbon stress-induced gluconeogenesis is terminated upon glucose availability, a process that involves the multiprotein E3 ligase, GID^SR4^, recruiting N-termini and catalyzing ubiquitylation of gluconeogenic enzymes. Here, genetics, biochemistry, and cryo electron microscopy define molecular underpinnings of ***g***lucose-***i***nduced ***d***egradation. Unexpectedly, carbon stress induces an inactive anticipatory complex (GID^Ant^), which awaits a glucose-induced substrate receptor to form the active GID^SR4^. Meanwhile, other environmental perturbations elicit production of an alternative substrate receptor assembling into a related E3 ligase complex. The intricate structure of GID^Ant^ enables anticipating and ultimately binding various N-degron targeting (i.e. “N-end rule”) substrate receptors, while the GID^SR4^ E3 forms a clamp-like structure juxtaposing substrate lysines with the ubiquitylation active site. The data reveal evolutionarily conserved GID complexes as a family of multisubunit E3 ubiquitin ligases responsive to extracellular stimuli.

## INTRODUCTION

Eukaryotes employ a plethora of mechanisms to cope with environmental perturbations. Much of our understanding of these processes comes from studies on the yeast *S. cerevisiae*, for example chaperone induction to enable protein folding during heat stress, kinase activation to control osmolarity, and glycolytic or gluconeogenic enzyme expression to switch metabolism. An emerging concept is that cells also have “anticipatory” programs whereby an altered growth condition not only triggers pathways rescuing cells from immediate dangers, but also expression of proteins that could be required for subsequent shifts in conditions (Mitchell et al., 2009; Tagkopoulos et al., 2008). If the anticipated perturbation does occur, cells can more rapidly adapt to the new environment through expression of yet other genes. For example, chaperones are induced at temperatures below those causing global misfolding, thereby increasing proteostasis capacity should a more severe later stress further compromise cellular protein folding (Klaips et al., 2014). Determination of protein fate by ubiquitylation is another major mechanism orchestrating homeostasis (Ciechanover, 2012; Varshavsky, 2012). Ubiquitylation depends on cellular signals directing E3 ligases to particular targets. Yet, our understanding of E3-dependent responses to environmental changes remains rudimentary. The questions of if and how E3 ligase structures play roles in cellular anticipation and responses to perturbations in the extracellular milieu are largely unexplored.

The ubiquitin (Ub) system has long been known to regulate yeast carbon catabolite repression (Zaman et al., 2008). While yeast growing on non-fermentable carbon sources (e.g. ethanol) require gluconeogenic production of glucose, this energetically costly pathway is futile and therefore terminated when sugars are available. This not only involves multifaceted transcriptional responses, but also *g*lucose-*i*nduced *d*egradation (Gid) of gluconeogenesis enzymes such as Fructose-1,6-bisphosphatase (Fbp1), malate dehydrogenase (Mdh2), and isocitrate lyase (Icl1) (Chiang and Schekman, 1991; Chiang and Chiang, 1998; Gancedo, 1998; Hoffman and Chiang, 1996; Schork et al., 1994a, b). The original Gid gene products defined by genetics and biochemistry include the E2 Ub conjugating enzyme Gid3 (hereafter referred to as Ubc8), the deubiquitylating enzyme Ubp14 (Gid6), and a GID complex loosely-defined by physical interactions of Gid1, Gid2, Gid4, Gid5, Gid7, Gid8, and Gid9 (Braun et al., 2011; Francis et al., 2013; Menssen et al., 2012; Regelmann et al., 2003; Santt et al., 2008; Schule et al., 2000). While the Gid2 and Gid9 subunits each harbor RING domains, the other subunits lack sequences associated with ubiquitylation (Braun et al., 2011; Francis et al., 2013; Menssen et al., 2012; Schule et al., 2000). A recent breakthrough in our understanding of the GID E3 came from its assignment as an N-degron-targeting E3 (Chen et al., 2017).

N-degron (formerly termed “N-end rule”) and C-degron (collectively referred to as “terminal degron”) E3s recognize substrate N- or C-termini and regulate vast biology (Varshavsky, 2019). Nonetheless, beyond knowledge of pathways creating, exposing, or cloaking substrate N- or C-degrons, and structures showing their recognition by E3 ligases, there is limited structural information explaining regulation of terminal degron E3s (Brower et al., 2013; Choi et al., 2010; Dong et al., 2018; Hu et al., 2005; Koren et al., 2018; Lin et al., 2018; Matta-Camacho et al., 2010; Rao et al., 2001; Rusnac et al., 2018; Shemorry et al., 2013; Szoradi et al., 2018; Timms et al., 2019; Varshavsky, 2011; Wang et al., 2008). Moreover, Fbp1, Mdh2, and Icl1 each harbor natively exposed GID E3-targeting N-terminal prolines essential for their degradation (Hammerle et al., 1998). The question of how their ubiquitylation could be regulated was answered by discovery that glucose availability determines expression of Gid4 (Menssen et al., 2018; Santt et al., 2008), which serves as a substrate receptor for the GID E3 by binding to an N-terminal proline (Chen et al., 2017; Dong et al., 2018). A crystal structure of peptide-bound human Gid4 showed the basis for N-terminal proline recognition (Dong et al., 2018). Although the mammalian GID E3 does not appear to regulate gluconeogenic enzymes (Lampert et al., 2018), and its N-degron substrates remain to be identified, numerous studies suggest it may also act as a central component in cell fate determination essential for some developmental pathways (Han et al., 2016; Javan et al., 2018; Liu and Pfirrmann, 2019; Nguyen et al., 2017; Pfirrmann et al., 2015; Soni et al., 2006)

Here we reveal molecular mechanisms underlying assembly and activity of the largely mysterious GID E3, and provide general insight into ubiquitylation by the large cohort of terminal-degron E3s, and by those catalyzing ubiquitylation via heterodimeric RING-RING domains. Unexpectedly, our results also reveal mechanisms of stress anticipation and resolution through assembly of an E3 ligase, and that GID is not a singular complex. GID comprises a family of multisubunit E3s regulated through assembly with interchangeable N-degron-binding substrate receptors induced by distinct environmental perturbations.

## RESULTS

### Carbon-source dependent anticipatory versus activated GID E3 ligase assemblies

As a prelude for developing and validating a recombinant system, we investigated properties of endogenous Gid proteins. The potential of Gid proteins to stably coassemble with each other *in vivo* was examined using a suite of yeast strains, each harboring a Gid gene tagged at its endogenous locus and validated for activity. Yeast were grown in various carbon sources known to determine GID E3 ligase activity (Oh et al., 2017; Regelmann et al., 2003), and lysates were subjected to sucrose gradient fractionation (Figure 1A).

**Figure 1.**
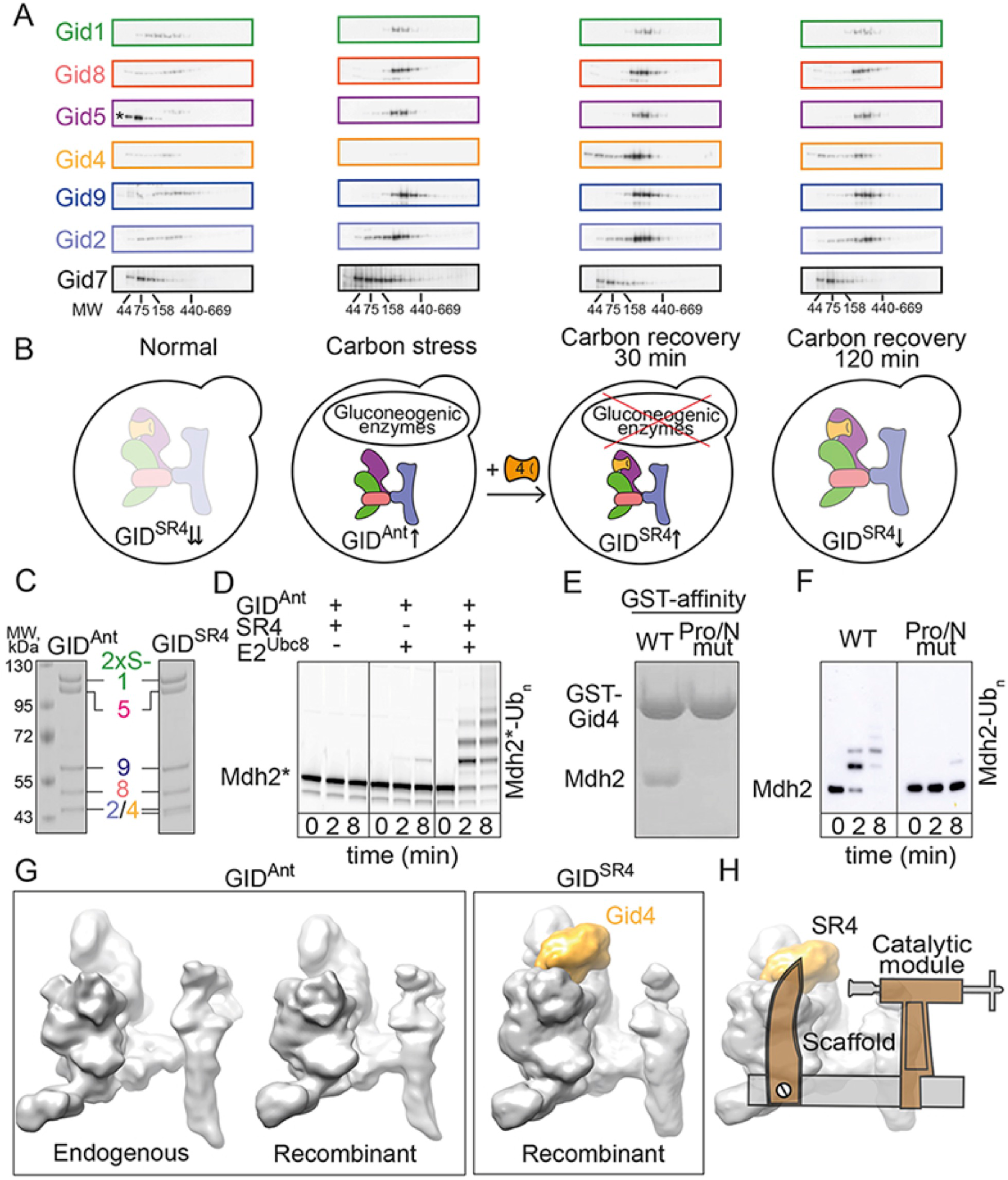
Regulation and reconstitution of yeast GID E3 ligase complexes. A. Sucrose density gradient fractionation of *S. cerevisiae* lysates from cells harvested from four conditions: stationary growth in glucose-rich medium (normal), ethanol (carbon stress), switch to glucose-rich medium for 30 and 120 minutes (carbon recovery 30 min and 120 min, respectively). Gid subunits tagged at their endogenous loci were visualized by western blotting. Asterisk indicates a non-specific anti-Flag interaction. B. Cartoons representing GID assemblies in different environmental conditions, based on migration patterns of subunits in sucrose density gradients. C. Coomassie-stained SDS-PAGE of recombinant GID^Ant^ and GID^SR4^. D. Fluorescent scan examining ubiquitylation of fluorescently-labelled Pro/N-degron substrate Mdh2 (Mdh2*). Assays test dependence on E2 (Ubc8) and substrate receptor (Gid4). Note: GID^Ant^ + Gid4 = GID^SR4^ E. Role of substrate N-terminal Pro, tested with WT Mdh2 or N-terminal Pro-to-Ser mutant, in binding GST-tagged substrate receptor Gid4 *in vitro*. F. Ubiquitylation of WT, or N-terminal Pro-to-Ser mutant, Mdh2-His_6_ visualized by western blot with anti-His_6_ tag antibody. G. 9 Å resolution cryo EM reconstructions of endogenous and recombinant GID^Ant^ and recombinant GID^SR4^. Density attributed to substrate receptor Gid4 is yellow. H. Clamp-like structure of GID^SR4^ assembled from Substrate Receptor Gid4 (SR4), Scaffold, and Catalytic modules.

Migration of Gid subunits, and their relative levels in the four conditions (Figure 1A, S1), led to three major conclusions (Figure 1B). First, in carbon recovery conditions that prompt degradation of gluconeogenesis enzymes, Gid1, Gid8, Gid5, Gid4, Gid9 and Gid2 comigrate, suggesting these subunits form a minimal stable E3 ligase including the ***<u>s</u>***ubstrate ***<u>r</u>***eceptor Gid***<u>4</u>***, that we term GID^SR4^. Second, as expected, the relative level of Gid4 is highest during carbon recovery, in agreement with Gid4 expression being the glucose-regulated switch determining E3 activity (Menssen et al., 2018; Santt et al., 2008). Finally and unexpectedly, during carbon stress, the levels of all GID^SR4^ subunits except Gid4 increase, and they comigrate in the density gradients. This suggests that during energetically expensive growth on a non-fermentable carbon source, a seemingly unnecessary, inactive complex containing most Gid proteins is produced. This finding can be rationalized by the emerging concept of “anticipatory” programs preparing for a later shift in conditions. Thus, we term the complex containing Gid1, Gid2, Gid5, Gid8 and Gid9 “GID^Ant^”, surmising that when produced during carbon stress GID^Ant^ would be benign toward gluconeogenic enzymes, but ready and primed for a potential later shift into glucose-containing media, which in turn would rapidly induce Gid4 expression and assembly of the active GID^SR4^ E3 ligase. Although there may be settings when GID^SR4^ and GID^Ant^ further include Gid7 *in vivo*, at this point the role of Gid7 remains unknown. Gid7 may bind to a subset of GID complexes, additional factors may contribute to its binding, the interaction may be transient or low affinity, or Gid7 may play alternative roles in regulation.

To mechanistically define regulation, we generated recombinant GID^Ant^ and GID^SR4^ complexes (Figure 1C) that reconstitute known GID features. First, together with the E2 enzyme Ubc8, GID^SR4^, but not GID^Ant^, catalyzed robust polyubiquitylation of a recombinant gluconeogenic enzyme substrate, Mdh2 (Figure 1D). Second, in accordance with impaired degradation of a Gid substrate upon overexpressing a dominant-negative Ub K48R mutant *in vivo* (Schork et al., 1995), we found that in the context of otherwise lysineless Ub only K48 supported substantial polyubiquitylation by our recombinant system (Figure S1D). Third, the N-terminal Pro of Mdh2 was required for its binding to Gid4 and ubiquitylation by GID^SR4^ (Figure 1E, 1F).

3D reconstructions at 9 Å resolution obtained by cryo electron microscopy (cryo EM) further validated our recombinant system. Comparing the EM maps for recombinant GID^Ant^ and that purified from yeast cultured in carbon stress conditions revealed a common overall architecture (Figure 1G). Thus, it appears that the native GID^Ant^ purified from yeast - at least in terms of subunits overtly visible by cryo EM at this resolution - is indeed a complex of Gid1, Gid2, Gid5, Gid8 and Gid9.

Prominent additional density correlating with the presence of the substrate receptor subunit, Gid4, was readily visible in the map of recombinant GID^SR4^ (Figure 1G). The overall structure of the GID^SR4^ E3 resembles a clamp, with Gid4 corresponding to one jaw (Figure 1H). A high-resolution structure showed the substrate receptor linked via a scaffold to a catalytic module as described below.

### Modular GID E3 ligase assembly

Refinement of the cryo EM data for GID^SR4^ yielded a 3D reconstruction at 3.8 Å resolution (Tables 1, 2, S1, S2, Figure S2-S6). Atomic coordinates for Gid4, Gid5, Gid8, and much of Gid1 and Gid9, were generated by a combinatorial approach involving cryo EM maps of many variant complexes, and automated and manual model building (Figure S2-S6).

**Table 1.**
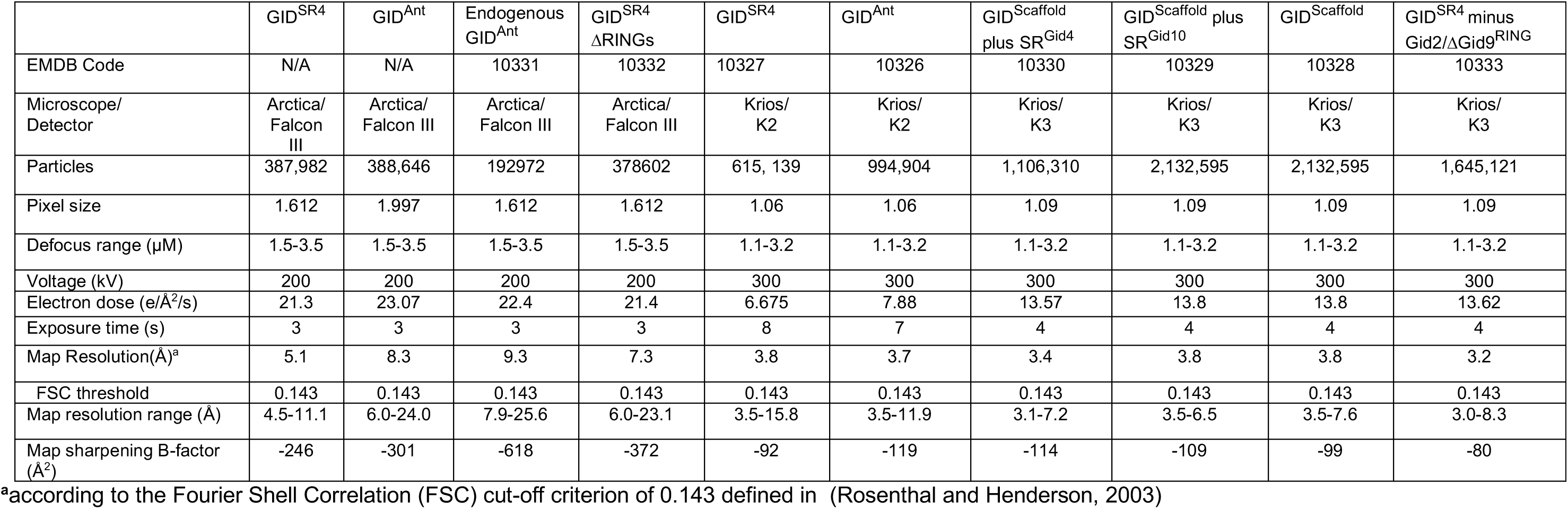
Cryo-EM Data Collection, 3D Reconstruction and Map Refinement.

| | GID<sup>SR4</sup> | GID<sup>Ant</sup> | Endogenous<br>GID<sup>Ant</sup> | GID<sup>SR4</sup><br>ΔRINGs | GID<sup>SR4</sup> | GID<sup>Ant</sup> | GID<sup>Scaffold</sup><br>plus SR<sup>Gid4</sup> | GID<sup>Scaffold</sup> plus<br>SR<sup>Gid10</sup> | GID<sup>Scaffold</sup> | GID<sup>SR4</sup> minus<br>Gid2/ΔGid9<sup>RING</sup> |
| --- | --- | --- | --- | --- | --- | --- | --- | --- | --- | --- |
| EMDB Code | N/A | N/A | 10331 | 10332 | 10327 | 10326 | 10330 | 10329 | 10328 | 10333 |
| Microscope/<br>Detector | Arctica/<br>Falcon<br>III | Arctica/<br>Falcon III | Arctica/<br>Falcon III | Arctica/<br>Falcon III | Krios/<br>K2 | Krios/<br>K2 | Krios/<br>K3 | Krios/<br>K3 | Krios/<br>K3 | Krios/<br>K3 |
| Particles | 387,982 | 388,646 | 192972 | 378602 | 615, 139 | 994,904 | 1,106,310 | 2,132,595 | 2,132,595 | 1,645,121 |
| Pixel size | 1.612 | 1.997 | 1.612 | 1.612 | 1.06 | 1.06 | 1.09 | 1.09 | 1.09 | 1.09 |
| Defocus range (μM) | 1.5-3.5 | 1.5-3.5 | 1.5-3.5 | 1.5-3.5 | 1.1-3.2 | 1.1-3.2 | 1.1-3.2 | 1.1-3.2 | 1.1-3.2 | 1.1-3.2 |
| Voltage (kV) | 200 | 200 | 200 | 200 | 300 | 300 | 300 | 300 | 300 | 300 |
| Electron dose (e/$\text{\AA}^2$/s) | 21.3 | 23.07 | 22.4 | 21.4 | 6.675 | 7.88 | 13.57 | 13.8 | 13.8 | 13.62 |
| Exposure time (s) | 3 | 3 | 3 | 3 | 8 | 7 | 4 | 4 | 4 | 4 |
| Map Resolution($\text{\AA}$)<sup>a</sup> | 5.1 | 8.3 | 9.3 | 7.3 | 3.8 | 3.7 | 3.4 | 3.8 | 3.8 | 3.2 |
| FSC threshold | 0.143 | 0.143 | 0.143 | 0.143 | 0.143 | 0.143 | 0.143 | 0.143 | 0.143 | 0.143 |
| Map resolution range ($\text{\AA}$) | 4.5-11.1 | 6.0-24.0 | 7.9-25.6 | 6.0-23.1 | 3.5-15.8 | 3.5-11.9 | 3.1-7.2 | 3.5-6.5 | 3.5-7.6 | 3.0-8.3 |
| Map sharpening B-factor<br>($\text{\AA}^2$) | -246 | -301 | -618 | -372 | -92 | -119 | -114 | -109 | -99 | -80 |

<sup>a</sup>according to the Fourier Shell Correlation (FSC) cut-off criterion of 0.143 defined in (Rosenthal and Henderson, 2003)

**Table 2.**
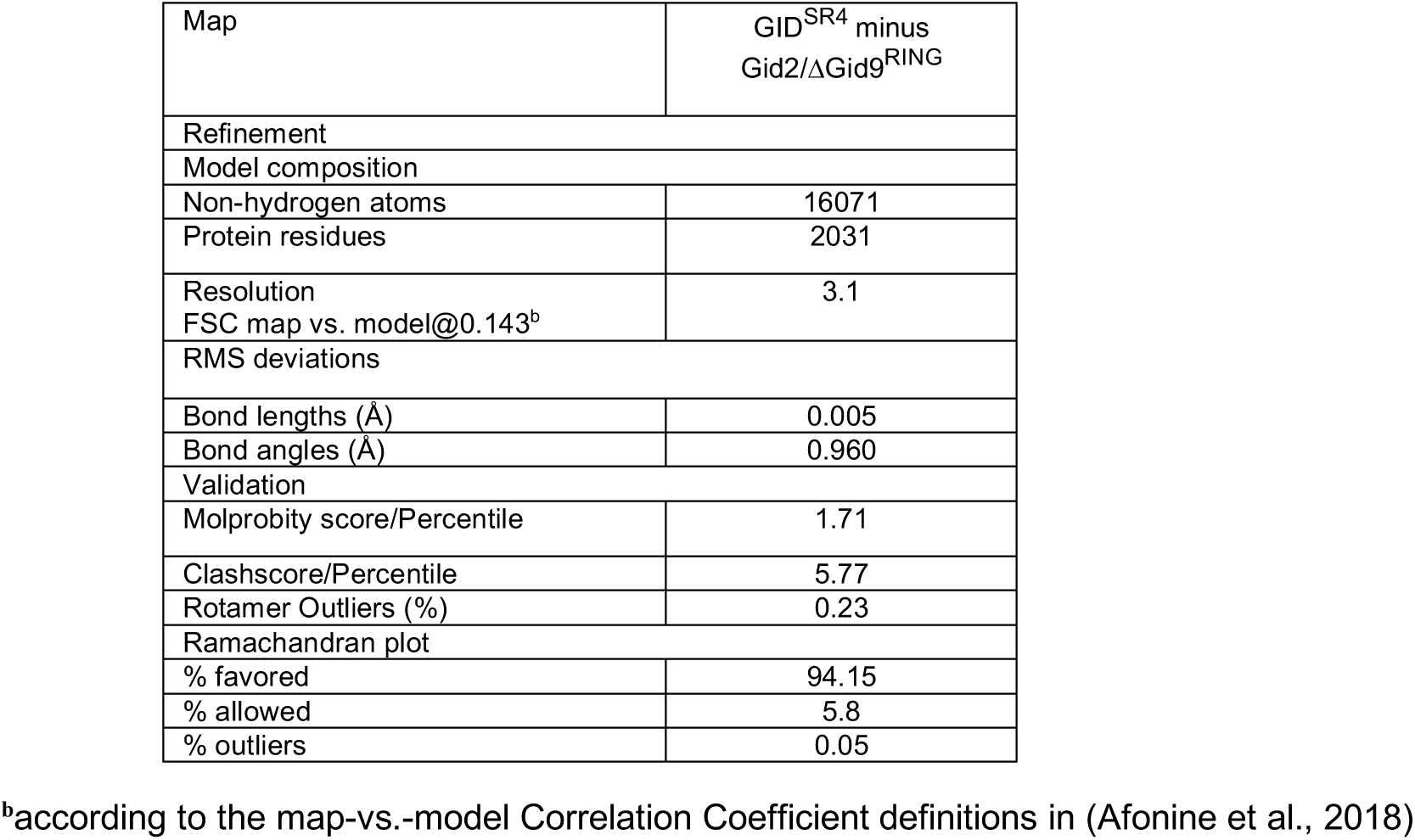
Model refinement and validation statistics.

Additional predicted domains from Gid1, Gid2 and Gid9 were approximately docked into lower resolution density (Figure S5B, S6A-B). The multidomain nature of Gid proteins enabled structure validation through: 1) testing effects of deleting specific subunits or domains on cryo EM reconstructions; 2) strong correlations upon superimposing structures of human Gid4 substrate-binding and Gid1 SPRY domains (1.0 and 0.73 RMSD, respectively) (Dong et al., 2018; Li et al., 2011); 3) visualizing predicted armadillo repeats in Gid5 and LisH-CTLH-CRA domains in Gid1, Gid8 and Gid9; and 4) concordance between the structure and effects of mutations observed in prior studies of GID E3 assembly *in vivo* (Braun et al., 2011; Menssen et al., 2012; Santt et al., 2008).

Overall, the EM data reveal that the GID E3 is organized around three structural and functional modules (Figure 2): the scaffold - Gid1, Gid5, and Gid8 tightly interacting in a manner that outwardly projects protein interaction domains from each subunit; the substrate receptor - Gid4; and the catalytic module - the Gid2–Gid9 subcomplex. Details of this assembly, and how it drives ubiquitylation of N-degron substrates, are described below.

**Figure 2.**
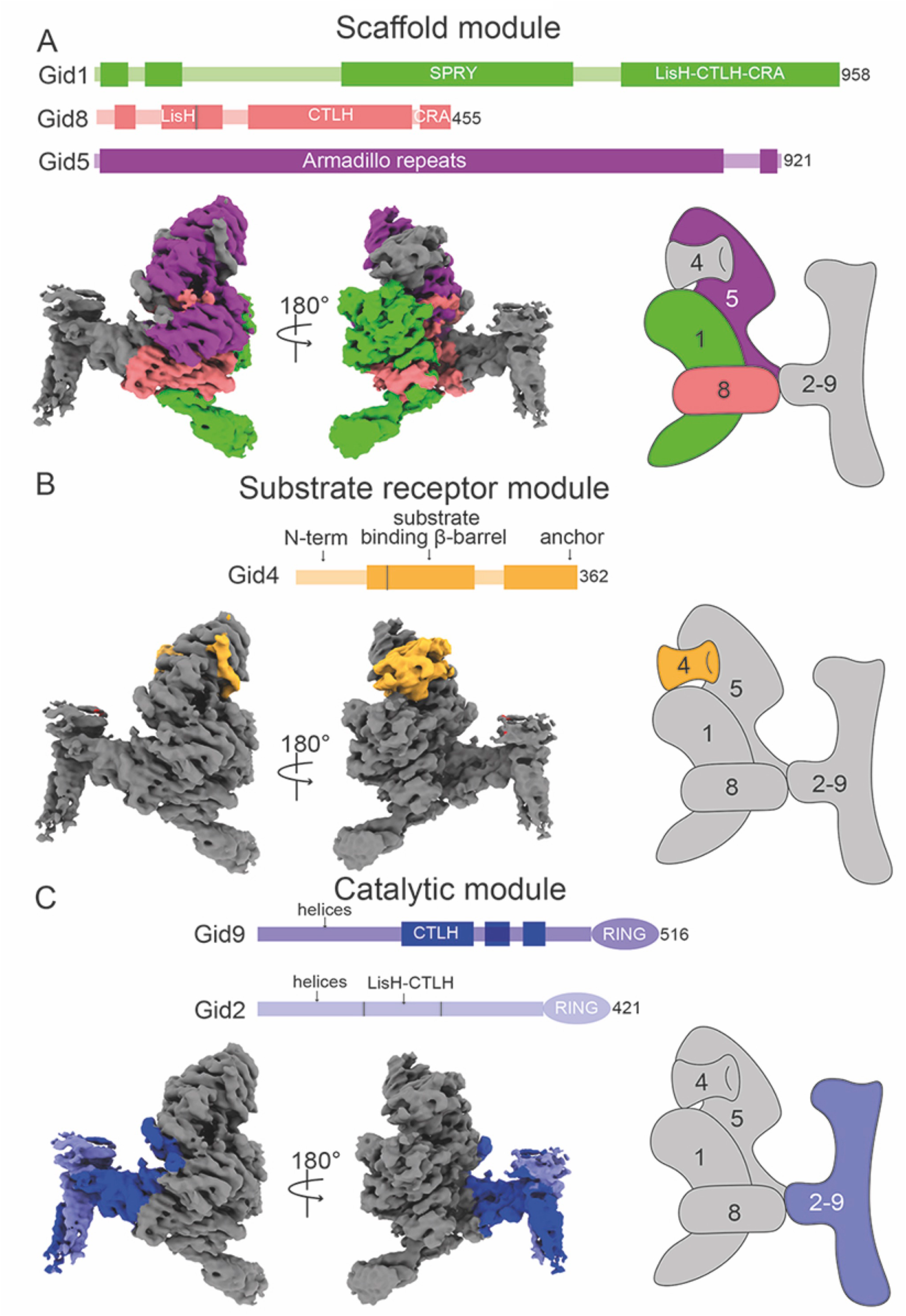
GID^SR4^ E3 ligase modular architecture. Each panel shows a different module as a domain schematic (top), 2 views of cryo EM density (bottom left) and a cartoon (bottom right). Subunits within a module are color-coded, with others in grey. Darker boxes in the domain schematic represent regions of the density map into which an atomic model was built. A. Scaffold module comprising Gid1 (green), Gid8 (salmon) and Gid5 (purple). B. Substrate receptor module consisting of Gid4 (orange) C. Catalytic module composed of Gid2 (light blue) and Gid9 (dark blue)

### The scaffold

The foundation of GID^SR4^ is an interdigitated assembly of Gid1, Gid8, and Gid5 (Figure 2A, S6C-F). Gid1 and Gid8 together form a heterodimeric trefoil-shaped structure. At the vertex, Gid1’s LisH and C-terminal segment of the CRA domain (LisH-CRA^C^), and adjacent elements, embrace paralogous regions from Gid8, rationalizing why Gid1 and Gid8 stabilize each other *in vivo* (Menssen et al., 2012). The three lobes of the trefoil are formed by 1) Gid1’s SPRY domain, 2) Gid1’s CTLH and N-terminal segment of the CRA domain (CRA^N^), and 3) Gid8’s CTLH-CRA^N^ domain and adjacent sequences (Figure S6C). The distal ends of the CTLH-CRA^N^ domains from both Gid1 and Gid8 radiate away from the core, while a continuous V-shaped surface between Gid1’s SPRY and Gid8’s CTLH domain engages an extended complementary surface from Gid5. Gid5’s armadillo repeats stack in tandem in a continuous solenoid of roughly one and a half superhelical turns, with the N-terminal domain (NTD) filling the groove between Gid1 and Gid8, and a C-terminal domain (CTD) radiating outward (Figure 2A, S6C, S6E). The scaffold is further buttressed by loops from all three proteins extending distances up to ≈70 Å to engage each other.

A protein interaction domain from Gid5 recruits the substrate receptor Gid4, and Gid8 binds the catalytic module Gid2–Gid9 (Figure S6C, S6F). Weak density corresponding to Gid1’s CTLH domain also projects outward. Although the structural role of Gid1’s CTLH domain is presently unknown, we speculate it binds Gid7 based on its mutation specifically impairing this interaction *in vivo* (Menssen et al., 2012).

### Scaffold binding to substrate receptor Gid4 generates GID^SR4^

The substrate receptor – which recruits proteins for ubiquitylation – is an essential E3 ligase element. A prior structure showed that human Gid4’s substrate-binding domain is largely a β-barrel, with a funnel-shaped opening at one end binding to short peptides via their N-terminal Pro (Figure 3A) (Dong et al., 2018). The structure of GID^SR4^ shows, in turn, how this substrate binding domain is incorporated into an active E3 ligase (Figure 2B, 3). Gid4’s C-terminal eight residues anchor the interaction, by extending into a channel in the concave surface of Gid5^CTD^ (Figure 3B, 3C). Here, successive Gid4 side chains protrude in opposite directions. Aromatic residues on one side fill pockets between Gid5 armadillo repeats. Those on the other side contribute to a composite Gid4/Gid5 interface with an aliphatic stripe across Gid4’s barrel. Indeed, mutation of key Gid4 and Gid5 contact residues impaired GID^SR4^-catalyzed substrate ubiquitylation *in vitro* (Figure 3D, 3E). *In vivo*, the Gid5 mutations substantially impaired degradation of the gluconeogenic enzyme Fbp1, as did mutation of Gid4’s C-terminal anchor. While individual conservative amino acid substitutions in Gid4’s substrate binding domain did not have a measurable effect, introduction of a bulky residue or multiple Ala mutations caused substantially impaired glucose-induced degradation of Fbp1 (Figure 3D-E).

**Figure 3.**
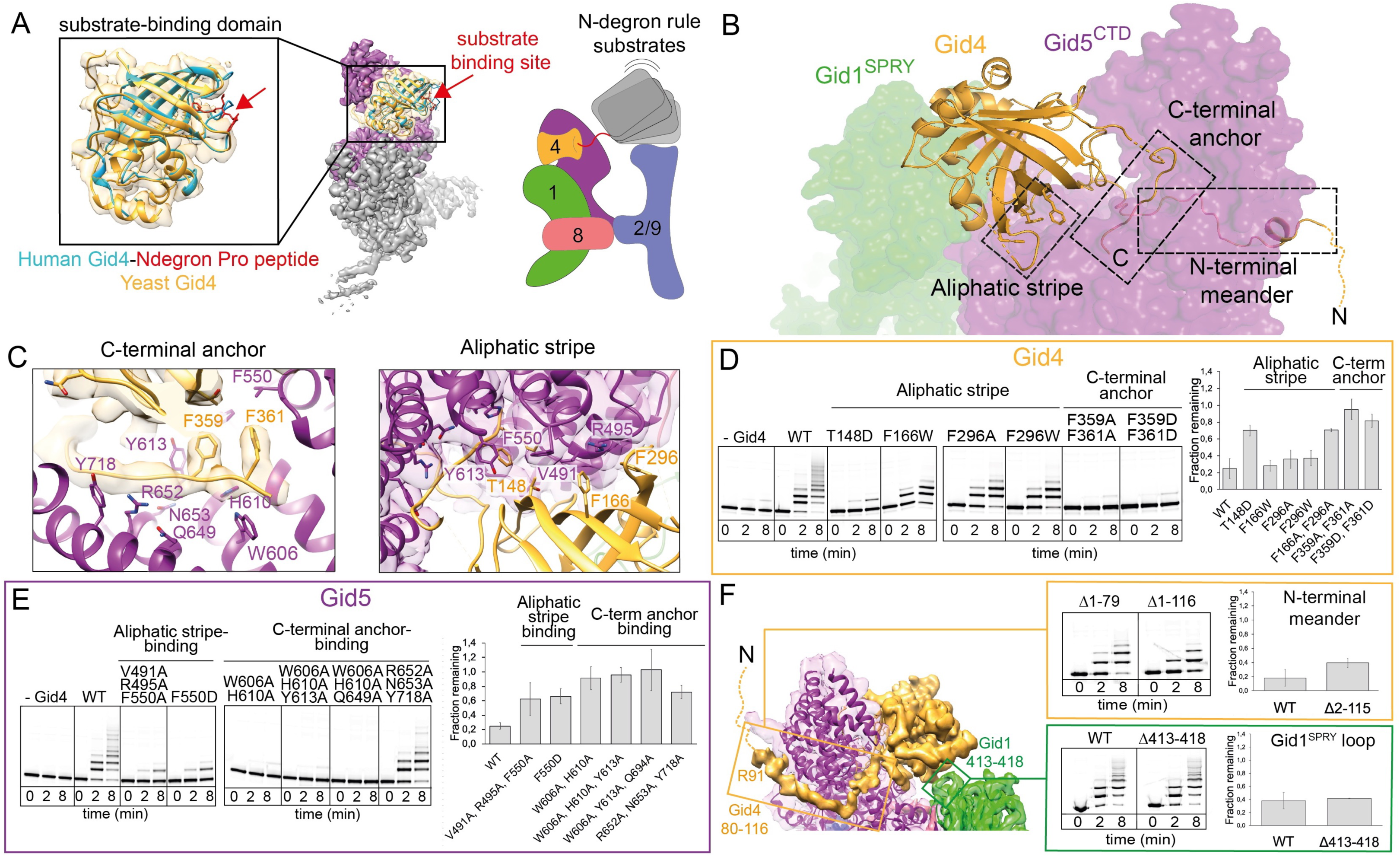
Formation of GID^SR4^ E3 ligase through incorporation of substrate receptor Gid4. A. Overlay of scaffold-bound *S. cerevisiae* Gid4 substrate-binding domain with crystal structure of human Gid4 bound to N-terminal Pro peptide (PDB 6CDC) (left), showing potential substrate binding site with a red arrow. Cartoon of N-degron substrate binding by GID^SR4^ (right). B. Overview of Gid4 elements binding to GID scaffold. Gid4 (yellow cartoon) binds Gid5^CTD^ (purple surface) via a C-terminal anchor (C=C-terminus), an aliphatic stripe and an N-terminal meander. Disordered residues connecting to N-terminus (N) shown as dotted line. C. Close-up of Gid4 (yellow) C-terminal anchor and aliphatic stripe interactions with Gid5 (purple). Residues mutated in D and E are represented as sticks. D. Assays testing importance of Gid4 residues in aliphatic stripe and C-terminal anchor on *in vitro* Mdh2 ubiquitylation and *in vivo* Fbp1 degradation (quantified as fraction from time 0 remaining after switching from carbon stress to carbon recovery). E. Assays testing importance of Gid5^CTD^ residues that interact with Gid4 aliphatic stripe and C-terminal anchor on *in vitro* Mdh2 ubiquitylation and *in vivo* Fbp1 degradation. F. Left, structure and EM density map depicting auxiliary interactions between Gid5^CTD^ and Gid4 N-terminal meander (residues 80-116) and between Gid1 SPRY domain loop (residues 413-418) and peripheral helical insertion in Gid4. Right, assays testing if these elements are not essential for *in vitro* ubiquitylation of Mdh2 and *in vivo* Fbp1 degradation.

Additionally, weaker EM density showed Gid4 residues 91-116, upstream of the substrate binding domain, meandering over 65 Å to loosely wrap around to the convex face of Gid5. Also a loop from Gid1’s SPRY domain contacts a peripheral helical portion of Gid4’s substrate-binding domain (Figure 3F). However, these residues are neither conserved nor essential for GID^SR4^ activity *in vitro* or *in vivo*, suggesting auxiliary roles (Figure 3F).

### Dynamic Gid5 CTD in anticipation of a substrate receptor

To understand the structure of the GID complex expressed during carbon stress, EM data for recombinant GID^Ant^ were refined to yield a 3D reconstruction at 3.7 Å resolution (Figure 4, S2, S4A, Tables 1, S1). Comparison with the map of GID^SR4^ showed a striking difference in the density corresponding to Gid5’s substrate-receptor binding CTD, which is blurred in GID^Ant^. The Gid5 armadillo repeats are visible, but poor density precluded refinement to high resolution (Figure 4A). Thus, we speculate that anticipation is manifested by conformational dynamics of the Gid5 CTD armadillo repeats prior to capturing and curling around a substrate receptor subsequently available upon change in environmental conditions.

**Figure 4.**
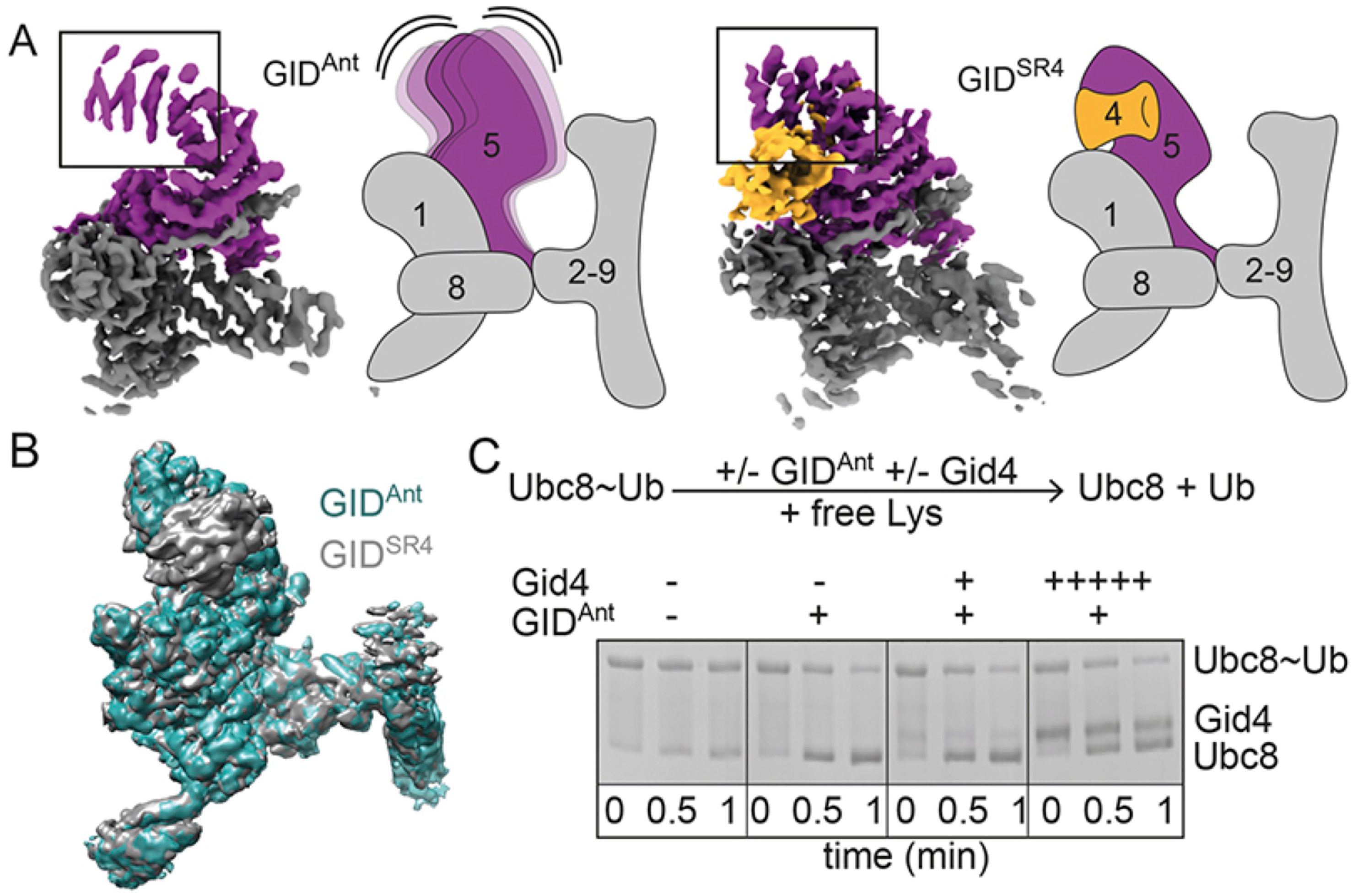
Structural anticipation by GID^Ant^. A. Cryo-EM maps and cartoons showing GID^Ant^ and GID^SR4^ with Gid5 purple and Gid4 yellow. Black boxes highlight relatively weaker Gid5^CTD^ density in GID^Ant^, which we presume represents conformational flexibility in the absence of substrate receptor. B. Superposition of cryo EM maps for GID^SR4^ (grey) and GID^Ant^ (aqua) at low contour. C. Assay testing substrate-independent E3 activity. First, the Ubc8∼Ub intermediate was generated enzymatically and this reaction was quenched. Next, free lysine was added. Reactivity probed by loss of Ubc8∼Ub and appearance of Ub was tested without an E3, or with GID^Ant^ alone, or with addition of equimolar and 5x excess of Gid4 over E3.

Because GID^Ant^ and GID^SR4^ are structurally similar beyond Gid5’s CTD and its associated Gid4 (Figure 4B), we hypothesized that GID complexes may display intrinsic catalytic activity irrespective of ability to recruit substrate. To test this, we employed an assay that monitors substrate-independent activation of E2∼Ub intermediates (Petroski and Deshaies, 2005; Wenzel et al., 2011). First, the reactive Ubc8∼Ub intermediate (“∼” refers to thioester linkage) was generated enzymatically and this reaction was quenched. Next, lysine was added simultaneously with various versions of GID E3s. Ub transfer from Ubc8, presumably to unanchored lysine, was monitored by both disappearance of Ubc8∼Ub and appearance of free Ubc8 in SDS-PAGE. While the Ubc8∼Ub intermediate was relatively stable on its own over time, GID^Ant^ stimulated its rapid discharge with little effect of titrating a version of Gid4 suitable for substrate recruitment (Figure 1D, 4C). Thus, GID^Ant^ is intrinsically competent at activating Ub transfer even without a recruited N-degron substrate or its receptor.

### A family of related GID E3s

The concept of a multiprotein E3 ligase that facultatively associates with a substrate receptor is conceptually reminiscent of cullin-RING and Anaphase-Promoting Complex E3 families. However, these E3s employ sets of interchangeable substrate receptors for distinct regulation (Alfieri et al., 2017; Lydeard et al., 2013; Watson et al., 2019). Thus, we hypothesized that other GID substrate receptor(s) may exist, and identified the ORF YGR066C as encoding a protein displaying homology to Gid4, including the Gid5-binding hydrophobic stripe and C-terminal anchor (Figure S4C). While our manuscript was under consideration, YGR066C was published as a GID E3 substrate receptor, and renamed “Gid10” (Melnykov et al., 2019). We have adopted this nomenclature, and had already independently performed several experiments suggesting that Gid10 is a bona fide alternative substrate receptor for a GID E3. First, bacterially expressed Gid10 binds our recombinant GID^Ant^ (Figure 5A). Second, Gid10 confers onto GID^Ant^ *in vitro* ubiquitylation activity toward an N-degron substrate, albeit with far lower efficiency than Gid4 (Figure 5B). Third, a 3.8 Å resolution cryo EM reconstruction of Gid10 bound to the scaffold module showed an overall similar structure to the Gid4-bound complex, including homologous placement of Gid10’s C-terminal anchor and a β-barrel domain poised to bind N-degron substrates (Figure 5C-D, S2, S4B, Table 1, S1). Indeed, deletion of Gid10’s C-terminal anchor abrogates Gid10-dependent ubiquitylation of the recombinant substrate Mdh2 (Figure 5B).

**Figure 5.**
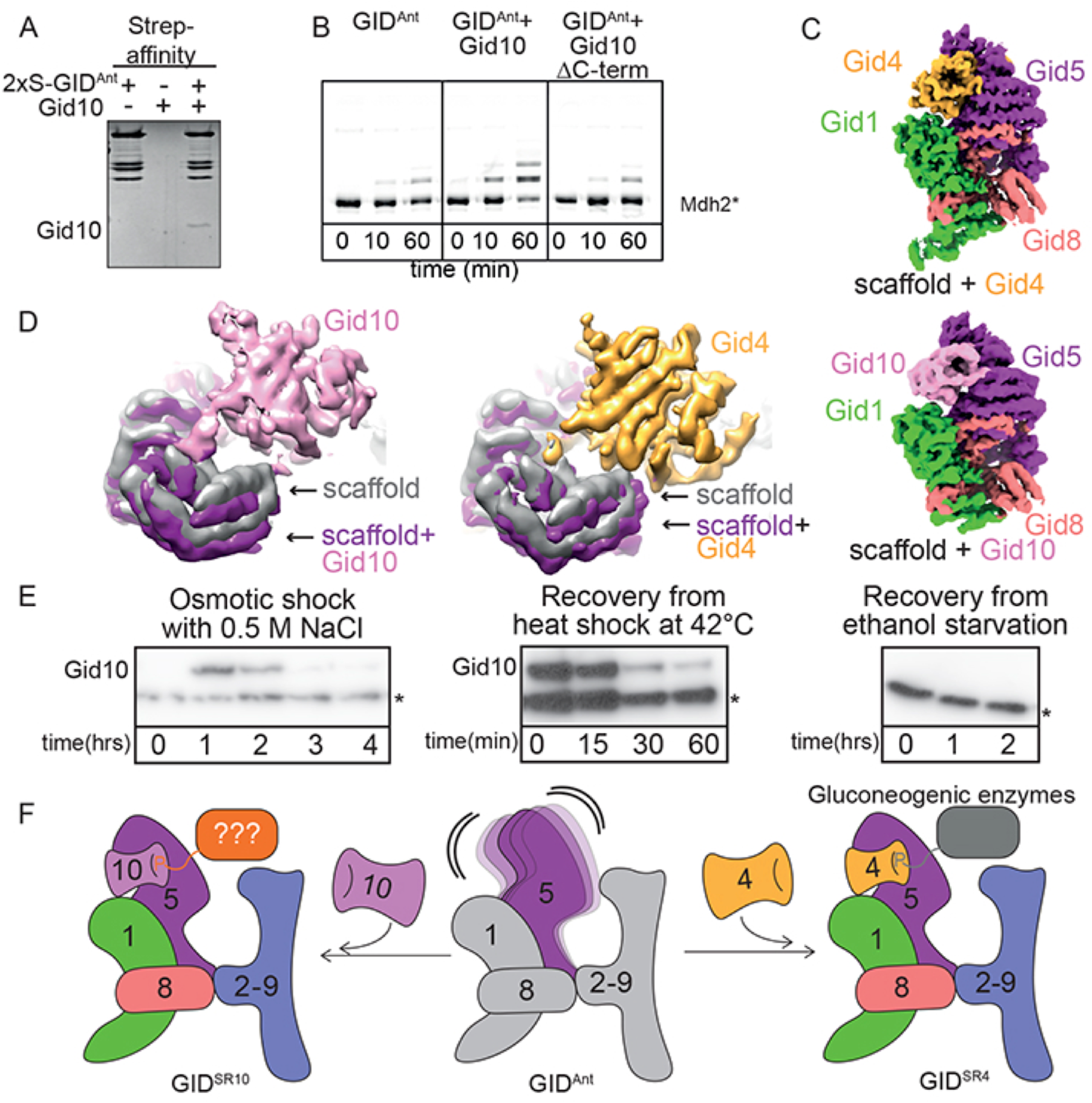
A family of multisubunit GID E3s with swappable substrate-receptors. A. Streptactin pull-down of GID^Ant^ testing binding of Gid10. B. Ubiquitylation assay testing potential of Gid10 to act as a substrate receptor for the GID^SR4^ substrate Mdh2. C. Cryo-EM maps of Gid4- and Gid10-bound GID scaffold. D. Close-up views of overlays of maps of GID scaffold alone and bound to Gid10 or Gid4. E. Western blots showing expression of Gid10, tagged at the endogenous locus, under different environmental conditions. Asterisk indicates a protein interacting non-specifically with anti-Flag antibodies. F. Model for family of GID E3s with interchangeable substrate receptors.

Comparing EM maps with the two substrate receptors in detail shows potential for slightly different placement of Gid10 and Gid4 relative to the scaffold (Figure 5C, 5D). This raises the possibility that orientation of substrate-binding domains may underlie mechanisms regulating substrate degradation under different cellular conditions.

Although deletion of the Gid10 gene in yeast did not impact degradation of known Gid substrates after carbon source switching (data not shown and (Melnykov et al., 2019)), prior transcriptomics, along with our analyses of protein levels, do not imply Gid10 expression under these conditions. Rather, Gid10 mRNA is expressed during various stresses, including high salinity and heat shock (Gasch et al., 2000; Wanichthanarak et al., 2014). Indeed, we observed Gid10 protein induction under these conditions, presumably leading to its incorporation into an alternative GID^SR10^ E3 complex (Figure 5E, 5F and (Melnykov et al., 2019).

### Embedding of a RING–RING catalytic module within multisubunit E3 ligase

Most E3 ligases depend on one or more RING domains binding to the E2 and the Ub in a thioester-linked E2∼Ub intermediate, thereby stabilizing a “closed conformation” that activates discharge of Ub from the E2 active site (Dou et al., 2012; Plechanovova et al., 2012; Pruneda et al., 2012). Thus, we sought to identify the structural locations and functional roles of the Gid2 and Gid9 RINGs. Having already placed the scaffold and substrate receptor modules, we attributed the remaining density to the catalytic module. This adopts a T-shaped structure, where the base of the T connects the catalytic domain to the scaffold (Figure 2, 6A, 6B, S6). Here, Gid9’s CTLH-CRA^N^ domain heterodimerizes with that from Gid8 in a manner resembling a pillar affixed to a base.

**Figure 6.**
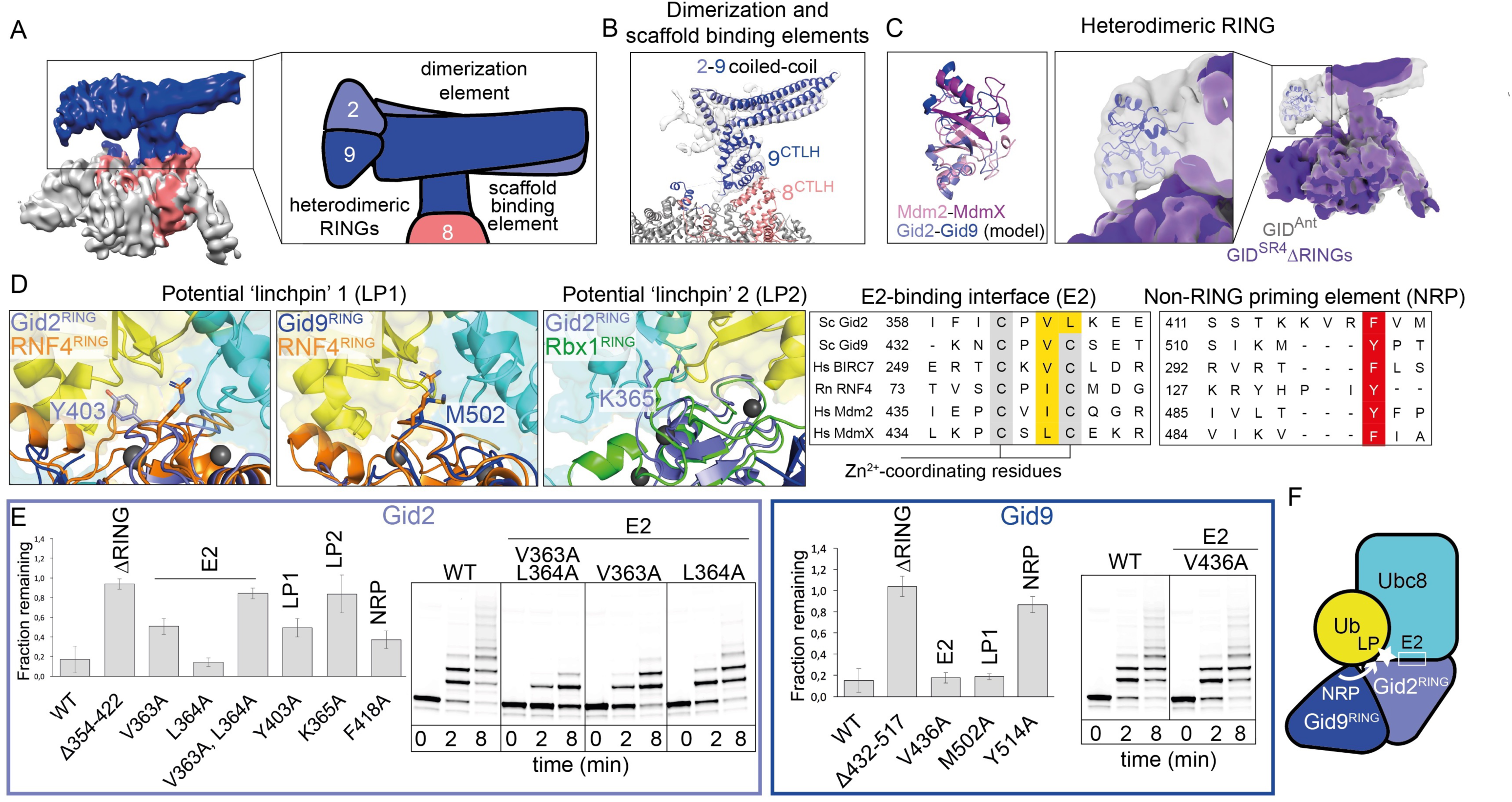
GID catalytic module. A. Left, T-shaped Gid2–Gid9 catalytic module in low contour EM map of GID^Ant^. Right, catalytic module elements shown in cartoon: scaffold-binding domain interacting with Gid8 (salmon), dimerization region, and heterodimeric RINGs. B. Homology models of catalytic module elements fitted into EM map generated by focused refinement and signal subtraction. The atomic models of Gid8 and Gid9 CTLH-CRA^N^ domains are shown, as is an approximated coiled-coil docked in additional density. C. The Gid2-Gid9 RING-RING domain was modeled in triangular density at the tip of the T-shaped catalytic module, as follows: 1) RING–RING domain was generated by superimposing homology models of Gid2 and Gid9 RINGs onto MDM2–MDMX structure (PDB 2VJE) (left). 2) Model of Gid2–Gid9 RING–RING domain was docked into map of GID^Ant^ (center). 3) Density attributed to the Gid2–Gid9 RING–RING domain was not visible in EM map of GID^SR4^ with the RINGs deleted (solid violet map, right). D. Left, candidate Gid2 and Gid9 RING ‘linchpins’ (LP) identified by superimposing their homology models with crystal structures of RNF4 (PDB 4AP4) and RBX1 (PDB 4P5O) bearing linchpin residues R181 and R46, respectively. Corresponding Gid2 and Gid9 residues are showed as sticks. Right, sequence alignments of Gid2 and Gid9 with well-characterized RING domains identified potential E2-binding (E2, yellow) and non-RING priming element (NRP, red) residues. E. Assays testing effects of Gid2 and Gid9 mutations on GID E3 activity, Fbp1 degradation *in vivo* and Mdh2 ubiquitylation *in vitro*. F. Cartoon summarizing model for Ubc8∼Ub activation by Gid2–Gid9 RING–RING domains based on mutational analysis shown in E.

The top of the T appears to comprise multiple heterodimeric Gid2–Gid9 subdomains. The relatively poor resolution of this region may suggest mobility of the Gid2–Gid9 subdomains with respect to each other, and relative to the scaffold. Although it was not possible to determine which elements derive from Gid2 or Gid9, the density was sufficiently visible at low contour to approximately localize predicted domains (Figure 6B, 6C, S5B, S6B). One side of the top of the T is a 4-stranded coiled coil, which we speculate corresponds to helices predicted at the N-termini of Gid2 and Gid9 (Kelley et al., 2015).

Most importantly, the structure of the catalytic core appears to place the Gid2 and Gid9 RING domains in a canonical RING–RING dimer assembly in the clamp-like structure of GID^SR4^, forming the second “jaw” that faces Gid4 (Figure 1H, 6C). We arrived at this conclusion after considering that the remainder of the T-structure consists of two subdomains, and then roughly attributing the unassigned Gid2–Gid9 features. The subdomain at the extreme edge of the complex can be fitted with a homology model of the Gid2 and Gid9 RING domains superimposed on a canonical RING–RING dimer assembly found in many E3 ligases (Kelley et al., 2015). Notably, the notion that the RINGs heterodimerize is consistent with prior mutations of zinc ligands within either protein, which presumably lead to RING misfolding, decreasing Gid2–Gid9 interactions and eliminating glucose-induced substrate degradation *in vivo* (Braun et al., 2011; Regelmann et al., 2003).

To validate the locations of the RINGs, we examined mutant versions of GID^SR4^ lacking these domains by cryo EM. Refinement of the data led to two major classes. One, indeed, showed selective elimination of the density we attribute to a Gid2–Gid9 RING–RING dimer, while the second class superimposed with the map obtained for a sample lacking the entire Gid2 subunit as well as the Gid9 RING domain (Table S1). This is consistent with heterodimeric assembly of the RING domains contributing to Gid2 incorporation into a GID E3 *in vivo* (Braun et al., 2011). Lastly, we speculate that the remaining density, at the T-junction corresponds to a heterodimeric assembly comprising the LisH-CRA^C^ domains from Gid2 and Gid9, and/or the ensuing CTLH domain from Gid2, which would match the size of this subdomain (Figure S6B). Moreover, this hypothesis is consistent with the relative orientation of Gid9’s CTLH-CRA^N^ domain, which is inserted between the LisH and CRA^C^ elements in the sequence of Gid9.

### Model of the catalytic center suggests the heterodimeric RING activates a single Ubc8∼Ub facing substrate

As a first step toward structurally modeling GID^SR4^-catalyzed ubiquitylation, each RING domain docked into the EM density was superimposed with a prior structure of an isolated RING–E2∼Ub complex (Dou et al., 2012; Plechanovova et al., 2012; Pruneda et al., 2012), and then the docked E2 was replaced with Ubc8. Even with uncertain position of the Gid2– Gid9 RING–RING dimer, the structural modeling suggested that only one of the two RING domains would place Ubc8 to face the Gid5-bound substrate receptor.

To test if the Gid2 and/or Gid9 RING primarily binds Ubc8 or plays a supporting role in activating the Ubc8∼Ub intermediate, residues were selected for mutation based on homology to three hallmark elements: 1) a hydrophobic surface that binds E2 loops conserved in Ubc8; 2) potential “linchpin” residues, which can be located on either side of the domain, but irrespective of location insert between the E2 and its thioester-linked Ub to stabilize the noncovalent interface between them; and 3) a non-RING priming element flanking a RING sequence that functions *in trans* to allosterically stabilize the closed conformation of the E2∼Ub intermediate bound primarily to the opposite RING in a dimer (Figure 6D) (Brown et al., 2014; Dou et al., 2012; Kelley et al., 2015; Plechanovova et al., 2012; Pruneda et al., 2012; Scott et al., 2014; Zheng et al., 2000). Effects on GID^SR4^ E3 ligase activity *in vivo* were tested by introducing mutations into tagged versions of Gid2 and Gid9 expressed from their endogenous loci (Figure 6E). Effects of point mutations in predicted E2-binding and linchpin residues of Gid2 mirrored effects of wholesale deletion of Gid2’s RING domain on glucose-induced degradation of Fbp1, while there was a relatively minimal effect of mutating Gid2’s candidate non-RING priming element. The crucial role for the Gid2 RING’s E2 binding site was also confirmed for GID^SR4^ E3 ligase activity *in vitro*. In contrast, the opposite pattern was observed for the Gid9 mutants, where only the candidate non-RING priming element significantly abrogated activity. The results suggest that Gid2’s RING binds and activates the Ubc8∼Ub intermediate, assisted by a non-RING priming element from Gid9, to face the substrate receptor (Figure 6E-F).

### Model of GID^SR4^ ubiquitylating an N-degron gluconeogenic enzyme

The substrate binding site on Gid4 is ≈50 Å away from the modeled catalytic center. While the relatively weak EM density corresponding to the catalytic domain (Figure 6C) suggests flexibility, perhaps for conformational changes during catalysis, it is also possible that the large gap accommodates substrates of GID^SR4^, which are large oligomeric enzymes. To model ubiquitylation, the substrate Mdh2 was selected because: 1) robust *in vitro* ubiquitylation of bacterially-expressed Mdh2 demonstrated that post-translational modifications are not required for its N-degron-based substrate targeting (Figure 1D-F) and as an ≈80 kDa homodimer with 34 lysines, Mdh2 is the smallest and structurally most simplistic of known GID^SR4^ substrates (Figure 7). To place Mdh2, the N-terminal four residues were modeled based on the prior structure of the human Gid4 substrate-binding domain bound to a 4-mer peptide. Next, an Mdh2 model was manually rotated while roughly constraining the location of the N-terminal-most ordered residue (L14) proximal to the substrate-binding site on Gid4 (Dong et al., 2018; Kelley et al., 2015).

**Figure 7.**
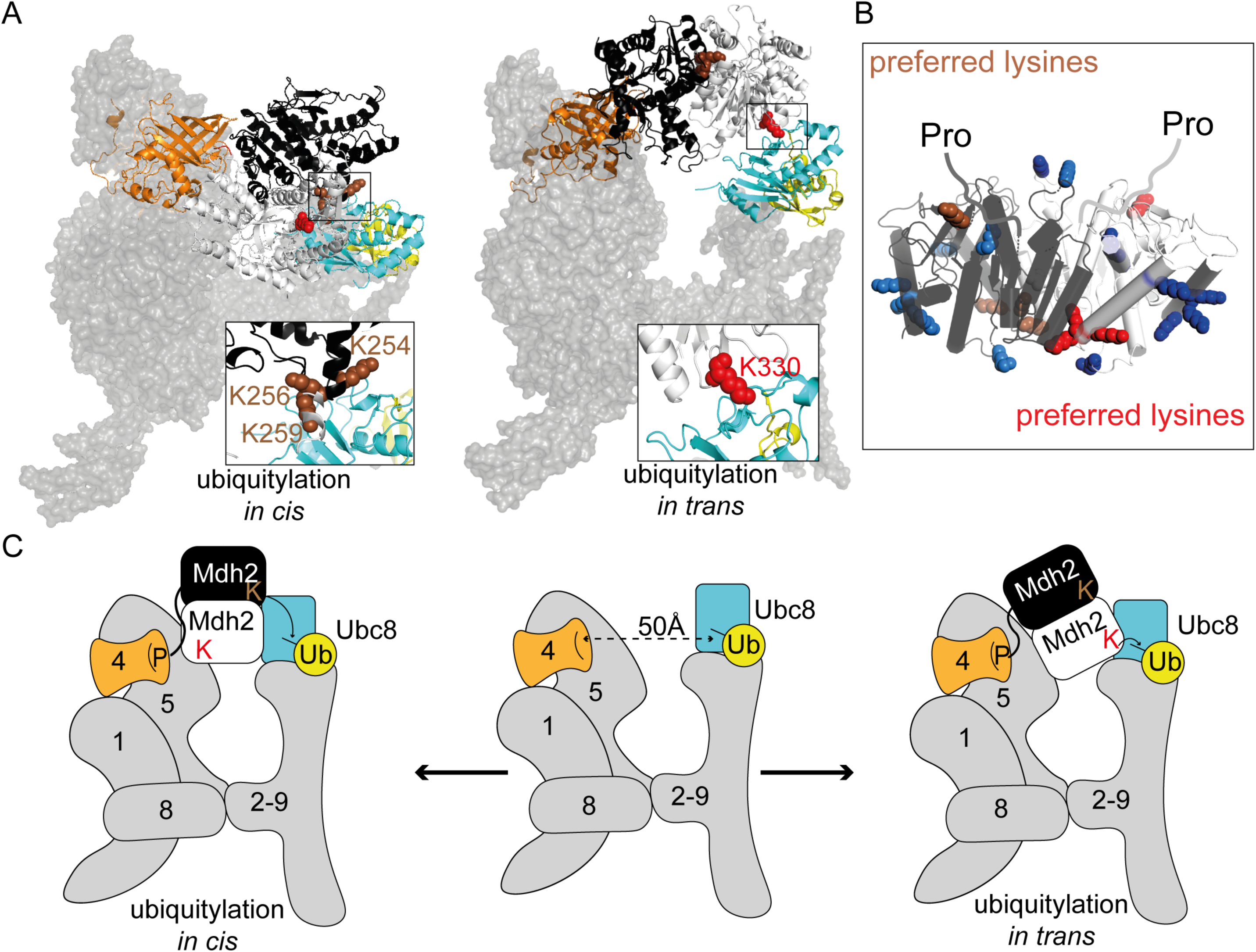
Model of GID^SR4^-catalyzed ubiquitylation of N-degron substrate Mdh2. A. Structural models for substrate ubiquitylation by GID^SR4^, with a homology model for Mdh2 (protomers in black and grey) placed with its N-terminal Pro binding Gid4 and candidate lysine targets in the active site of a modeled Ubc8∼Ub intermediate. B. Homology model of Mdh2 dimer showing preferred target lysines identified by mass spec in brown and red and other lysines in blue. C. Cartoon representing the structural models for GID^SR4^ ubiquitylation. The clamp-like structure enables multiple configurations for ubiquitylation of preferred lysines from a folded gluconeogenic enzyme substrate.

Overall, the model suggests that some, but not all, Mdh2 lysines would preferentially access the ubiquitylation active site (Figure 7A). To test this, we used mass spectrometry to map sites that are ubiquitylated *in vitro* (Figure S7). Notably, the top sites include a cluster of K254, K256, and K259, as well as K330, and to a lesser extent K360 and K361, for which the 10-residues between Mdh2’s N-terminus and globular domain would easily accommodate the ≈10, ≈20 and ≈15 Å required respectively for Mdh2 to simultaneously engage Gid4 and approach the Gid2 RING activated Ubc8∼Ub intermediate. Thus, while detailed knowledge awaits further structural studies, the EM data presented here enables generating a geometrically reasonable model for N-degron substrate ubiquitylation (Figure 7C).

## DISCUSSION

The cryo EM reconstructions reported here reveal E3 ligase assemblies that vary in response to extracellular stimuli (Figure 1, 4, 5, S1), molecular mechanisms underlying their regulation (Figure 1-6), and a framework for GID E3 ligase-dependent ubiquitylation (Figure 7). The structural data also provide broad insights into major families of E3 ligases, namely those recognizing terminal degrons and those displaying RING-RING catalytic domains. The modular multiprotein GID^SR4^ E3 assembly displays clamp-like properties, established by a central scaffold connecting the two jaws: a variable substrate receptor and the catalytic domain. The structure enables binding of a substrate’s unfolded N-terminus to Gid4, to direct lysines from a folded domain into the ubiquitylation active site. From the perspective of the other side of the complex, it seems that the RING–RING dimer is the culmination of an intricate heterodimeric Gid2–Gid9 assembly that activates the Ubc8∼Ub intermediate facing the N-degron substrate. Some other E3s, for example BRCA1–BARD1 or HDM2–HDMX, that contain heterodimeric RING–RING domains, may likewise rely on complex, interconnected assemblies to couple a single active site with a substrate for ubiquitylation.

While it has long been recognized that GID^SR4^ is part of the yeast response to environmental conditions via glucose-induced degradation of gluconeogenic enzymes (Santt et al., 2008), our data indicate that generation of a GID^Ant^ complex also occurs in response to an extracellular stimulus: carbon stress (Figure 1A). As GID^Ant^ would be inactive toward recruited substrates in the absence of a substrate receptor, we hypothesize that production of this complex allows cells to adapt more rapidly to potential later changes in the extracellular milieu. Our data raise the possibility that carbon stress may prepare cells for a potential return to nutrient-rich conditions. We also cannot rule out the possibility that GID^Ant^ could be coupled to a yet unknown substrate receptor to allow ubiquitylation of a distinct set of proteins during carbon stress.

We were puzzled by the apparently counterintuitive carbon stress-induced production of GID^Ant^ in anticipation of relief from starvation. It is conceptually appealing to envisage microbial anticipatory signaling cascades in terms of one stress serving as a signal for cells to cross-prepare for a looming new stress (Mitchell et al., 2009; Tagkopoulos et al., 2008). Our discovery that recombinant GID^Ant^ also binds the Gid4-like protein YGR066C/Gid10, which is induced under several distinct stress conditions (Figure 5 and (Melnykov et al., 2019)), offers a potential mechanism for how GID^Ant^ could act as a multi-faceted hub integrating responses to various extracellular stimuli. We speculate that carbon stress-induced production of GID^Ant^ may enable cells to prepare for ensuing osmotic stress or heat stress through the production of a Gid10-associated GID^SR10^ E3 ligase. It seems plausible that Gid10’s substrates could be regulators of glycerol or salt intake, protein synthesis, or general stress responses. Mechanistically, it seems likely that substrate selectivity will be influenced not only by protein expression changes under different metabolic conditions, but also by subtle differences in the β-barrel domains of Gid4 and Gid10, and their orientations relative to the scaffold (Figure 5). Future studies will be required to identify substrates of a GID^SR10^ E3, to visualize substrate ubiquitylation in action, and to understand cross-talk between GID^Ant^ assembly, association with multiple substrate receptors, and coupling responses to varying extracellular stimuli.

What does an E3 ligase in “anticipation” look like? Our cryo EM reconstruction of GID^Ant^ suggested motion of Gid5’s substrate receptor-binding CTD (Figure 4). Dynamic opening and closing of Gid5 could enable binding, release, and exchange of the substrate receptor. At this point, the molecular stimuli and structural mechanisms underlying substrate receptor dissociation or exchange remain unknown, although Gid4 turnover has been shown to depend on GID E3-dependent ubiquitylation (Menssen et al., 2018).

We speculate that in humans, a GID^Ant^ complex will parallel its yeast counterpart and act as a hub integrating various signals, presumably through many additional binding partners, in turn leading to cell fate determination. While binding to Gid4 likely generates a human GID^SR4^ E3 targeting substrates with N-terminal prolines (Dong et al., 2018), the functions of other partner proteins, including Gid7, remain elusive, and may regulate activity of GID^SR4^, or perhaps form alternative assemblies with GID^Ant^ or Gid subunits. Indeed, two Gid7 homologs, along with many other proteins, have been shown to co-precipitate with human Gid subunits (Boldt et al., 2016; Huttlin et al., 2017; Lampert et al., 2018). In addition, the human GID E3 ligase has been reported to ubiquitylate a substrate that does not bear an N-terminal proline (Lampert et al., 2018), despite the apparent requirement of a proline to bind human Gid4 (Dong et al., 2018). This could potentially be reconciled based on our discovery that the GID E3 ligase is not a singular complex but a family of E3 ligases with different substrate receptors (Figure 5). Additional human Gid subunits could substitute for Gid4, modulate substrate specificity, or localize the GID complex (Boldt et al., 2016; Lampert et al., 2018).

The concept of multiple GID E3 assemblies responding to different environmental stimuli is reminiscent of other multiprotein E3 ligases (e.g. cullin RING-ligases) and hubs such as mTOR that integrate signaling with various downstream functions required in certain cellular contexts (Gonzalez and Hall, 2017; Lydeard et al., 2013; Saxton and Sabatini, 2017). Regulation of these assemblies through interchangeable receptors provides a framework for investigating the GID family. Are there cellular exchange factors that promote swapping Gid4 and Gid10 (Pierce et al., 2013), or inhibitory factors (Duda et al., 2012; Lyapina et al., 2001)? Is GID regulated by modifications or metabolites (Gonzalez and Hall, 2017; Saxton and Sabatini, 2017)? Does substrate binding play a role in substrate-receptor selection (Emberley et al., 2012; Enchev et al., 2012)? And most curiously, are there other presently unknown substrate receptors? Although future studies will be required to unveil the molecular pathways and mechanisms underlying these complexities, the stunning structural intricacies of the seemingly simple yeast GID N-degron targeting system - now revealed more than 25 years since the discovery of glucose-induced degradation - provide a blueprint for understanding this important family of multisubunit E3 ligases.

## ACKNOWLEDGEMENTS

We thank J. Frye and support from ALSAC/St. Jude for cloning Gid ORFs; A. Varshavsky for plasmids for promoter reference assays in yeast; J. Kellermann for assistance in all work in Schulman lab; N. Nagaraj, V. Sanchez Caballero, N. Krombholz and A. Wehner for intact mass spec; I. Paron, C. Deiml and J. Mueller for mass spec maintenance; and M. Strauss, D. Scott, J.W. Harper and members of the Schulman lab for discussions. This was supported by Max Planck Society.

## AUTHOR CONTRIBUTIONS

SQ established recombinant GID and prepared cryo EM samples; SQ, DB, TS, JRP collected cryo EM data; SQ, JRP determined, refined, validated atomic models; CL, VB generated yeast strains; CL, VB, SQ performed yeast biochemistry; SQ, JC, DS performed in vitro biochemistry; OK, FH performed mass spec supervised by MM; SvG produced baculoviruses and insect cells expressed GID complexes; SQ, JC, DS, CL, BAS wrote the paper; AFA and BAS coordinated and supervised.

## METHODS DETAILS

### Yeast strains and growth conditions

All yeast strains used in this study are listed in Key Resource Table and are derivatives of BY4741. For strain construction, standard genetic techniques were employed (Janke et al., 2004; Knop et al., 1999; Storici and Resnick, 2006). All yeast strains were verified by DNA sequencing, western blotting for protein expression, and were shown to be competent for Fbp1 degradation (see below).

Unless otherwise specified, for assays described here, yeast strains were grown to OD_600_ of 1.0 in synthetic complete (SC) medium (0.17% yeast nitrogen base, 0.5% ammonium sulfate, 2% glucose, plus a mixture of amino acids). If strains were carrying a plasmid, the appropriate amino acids were omitted. Cells were then centrifuged at 1900xg for 3 minutes, washed once with SE medium (0.17% yeast nitrogen base, 0.5% ammonium sulfate, 2% ethanol, plus a mixture of amino acids), and then resuspended in fresh, pre-warmed SE media to an OD_600_ of 1.0. Cells were grown at 30°C for 19 hours, at which point they were harvested by centrifugation at 1,900xg for 3 minutes, and resuspended to an OD_600_ of 1.0 in fresh SC medium. At the indicated time points, cells were harvested by centrifugation at 11,200xg for 2 minutes, and flash frozen in liquid nitrogen for later analysis.

For growth under heat shock conditions, cells were grown in YPD at 30°C to mid-log phase and then shifted to 42°C for 30 minutes, before returning the cultures to 30°C growth. For growth under high salinity conditions, cells were grown to mid-log phase in YPD, then pelleted and resuspended in fresh YPD + 0.5 M NaCl and allow to grow at 30°C. At the indicated time points, an aliquot of cells was harvested by centrifugation.

### Fbp1 degradation assays

Fbp1 degradation assays were carried out using the promoter reference technique as previously described (Oh et al., 2017). Briefly, cells were first transformed with a plasmid co-expressing Fbp1 and a control protein (DHFR) from identical promoters containing an element that once transcribed binds tetracycline to inhibit translation. After growth for 19 hours in medium containing 2% ethanol, cells were resuspended to an OD_600_ of 1.0 in SD medium lacking the appropriate amino acids and containing 2% glucose and 0.5 mM tetracycline. At the indicated timepoints, 1 OD_600_ equivalent of cells were harvested. Cells were lysed by resuspension in 0.2 M NaOH followed by incubation on ice for 20 minutes, and then pelleted by centrifugation at 11,200xg for 2 minutes. The supernatant was removed and the cell pellet was resuspended in HU buffer containing 1X complete protease inhibitor tablets (Roche), heated at 70°C for 10 minutes, and then the resulting lysate was precleared by centrifugation for 5 minutes at 11,200xg. Samples were loaded on a 12% SDS-PAGE gel, followed by analysis by western blotting. Blots were imaged on a Typhoon scanner (GE Healthcare) and bands were quantified using ImageStudio software (Licor). At each time point, the amount of Fbp1 was normalized to the DHFR control protein.

### Protein digestion of in vitro ubiquitylation assays

Protein concentrations of the samples from *in vitro* ubiquitylation assays were measured by Bradford assay (BioRad). 4 μg of samples were 4-fold diluted in digestion buffer (1 M Urea in 50 mM Ammonium Bicarbonate, pH 8.0) followed by addition of TCEP and CAA to a final concentration of 10 mM and 40 mM, respectively, and incubated for 5 min at 45°C for reduction and alkylation. The samples were either digested using Trypsin (1:20 w/w, Sigma-Aldrich) alone, Trypsin (1:40 w/w)/GluC (1:40 w/w, BioLab) or Trypsin (1:40 w/w)/AspN (1:40 w/w, Promega) at 37°C overnight. In all cases, protease activity was quenched by acidification with Trifluoracetic acid to concentration of 1%.

### LC-MS/MS sample preparation

Acidified samples were loaded onto SDB-RPS StageTips, pre-equilibrated with 30% Methanol /1% Trifluoracetic acid and washed with 0.2% Trifluoracetic acid. StageTips were prepared by inserting two layers of SDB-RPS matrix (Empore) into a 200 μl pipette tip using an in-house prepared syringe device as described previously (Kulak et al., 2014). The StageTips were centrifuged at 1000xg. Loaded samples were sequentially washed with 0.2% Trifluoracetic acid and 2% Acetonitrile/0.2% Trifluoracetic acid, followed by elution with 1.25% NH_4_OH/80% Acetonitrile. Eluates were dried using a SpeedVac centrifuge (Eppendorf, Concentrator plus). Peptides were resuspended in buffer A* (2% Acetonitrile /0.1% Trifluoracetic acid) and briefly sonicated (Branson Ultrasonics) before LC/MS-MS analysis.

### LC-MS/MS Measurements

Peptide concentration was estimated by UV spectrometry and approximately 200 ng was loaded on a 50 cm reversed phase column (75 μm inner diameter, packed in house with ReproSil-Pur C18-AQ 1.9 μm resin [Dr. Maisch GmbH]). Column temperature was maintained at 60°C using a homemade column oven. Peptides were separated with a binary buffer system of buffer A (0.1% Formic acid (FA)) and buffer B (80% Acetonitrile plus 0.1% FA), at a flow rate of 300 nl/min. We used an EASY-nLC 1200 system (Thermo Fisher Scientific), which was directly coupled online with the mass spectrometer (Q Excative HF-X, Thermo Fisher Scientific) via a nano-electrospray source. Peptides were eluted with a gradient starting at 3% buffer B and stepwise increased to 8% in 8 min, 36% in 32 min, 45% in 4 min and 95% in 4 min. The mass spectrometer was operated in Top12 data-dependent mode (DDA) with a full scan range of 250-1350 m/z at 60,000 resolution with an automatic gain control (AGC) target of 3e6 and a maximum fill time of 20 ms. Precursor ions were isolated with a width of 1.4 m/z and fragmented by higher-energy collisional dissociation (HCD) with a normalized collision energy (NCE) of 28%. Fragment scans were performed at a resolution of 30,000, an AGC of 1e5 and a maximum injection time of 110 ms. Dynamic exclusion was enabled and set to 15 s.

### LC-MS/MS raw data processing

Raw MS data were searched against the UniProt Yeast FASTA using MaxQuant (version 1.6.2.10) (Cox and Mann, 2008; Cox et al., 2011) with a 1% FDR at peptide and protein level. Cysteine carbamidomethylation was set as fixed, protein N-Terminal acetylation, methionine oxidation and lysine diGly as variable modifications. The minimum peptide length was set to 7 amino acids, enzyme specificity was set to trypsin and two missed cleavages were allowed, permitting a maximum of 5 modifications per peptide. MS/MS spectra identifying ubiquitylated peptides of interest were obtained and exported using MaxQuant Viewer (Tyanova et al., 2015).

### Density Fractionation by sucrose gradients

At the indicated time points, 100 OD_600_ equivalents were harvested by centrifugation at 1900xg for 3 minutes. Cells were resuspended in lysis buffer containing 50 mM Tris-HCl pH 7.5, 150 mM NaCl, 1 mM MgCl_2_, 1% NP-40, and protease inhibitors (Roche), and lysed by glass bead lysis using a FastPrep-24 (MP Biomedicals). The resulting lysate was pre-cleared by centrifugation at 4,000xg for 10 minutes, and the supernatant was normalized by Bradford for total protein content, loaded onto a 5-40% sucrose gradient, and centrifuged at 34,300 rpm for 16 hours at 4°C. Each gradient was harvested into fourteen equal fractions, and run on SDS-PAGE, followed by analysis by western blotting with the appropriate antibody. Approximate molecular weights for fractions were determined using the protein standards provided with Gel Filtration Calibration Kit HMW (GE Healthcare). Briefly, 2 mg of each protein standard were resuspended in lysis buffer and run on a 5-40% gradient as described above.

### Plasmids preparation and Mutagenesis

Genes encoding GID subunits and Mdh2 substrate were originally amplified using *S. cerevisiae* BY4742 genomic DNA as a template. Upon initiating functional studies, we noticed that all sequences except Gid5 match those in the Saccharomyces Genome Database (SGD), where the Gid5 sequence corresponds to accession number NP_012247. The sequence of Gid5 used in the structure corresponds to accession number AJR43361 from the strain YJM1133. The difference is a single Gid5 Y758N residue substitution. We tested the functionality of the sequence used in the structure in initial assays probing GID^SR4^ activity toward Mdh2, and we confirmed by cryo EM that the overall structures of GID^SR4^ are similar with the two versions of Gid5. We converted to the SGD sequence for functional studies.

The genes encoding GID subunits were combined into one baculoviral expression vector with the biGBac method (Weissmann et al., 2016). All the plasmids used in this study are listed in the Key Resource Table.

The constructs for recombinant protein expression were generated by Gibson assembly method (Gibson et al., 2009) with a home-made Gibson reaction mix. To generate all the mutant versions of the constructs, QuickChange (Stratagene) protocol was applied. All coding sequences used for protein expression were entirely verified by sequencing.

### Protein expression and purification for cryo-EM

GID complexes and all the subcomplexes used for the single particle cryo EM analysis (GID^SR4^, GID^Ant^, GID^Scaffold^ plus substrate receptor Gid4 (SR^Gid4^), GID^Scaffold^ plus substrate receptor Gid10 (SR^Gid10^), GID^SR4^ minus Gid2/ΔGid9^RING^, GID^SR4^ ΔRINGs), were expressed in insect cell. For protein expression, Hi5 insect cells were transfected with recombinant baculovirus variants carrying the respective protein coding sequences and grown for 60 to 72 hours in EX-CELL 420 Serum-Free Medium at 27°C.

Cell pellets were resuspended in a lysis buffer containing 50 mM Tris-HCl pH 8.0, 200 mM NaCl, 5 mM DTT, 10 μg/ml leupeptin, 20 μg/ml aprotinin, 2 mM benzamidine, EDTA-free protease inhibitor tablet (1 tablet per 50 ml of the buffer) and 1 mM PMSF. The tagged complexes were purified from cell lysates by Strep-Tactin affinity chromatography by pulling on the Twin-Strep tag fused at the Gid1 N-terminus. Elutions were further purified by anion exchange chromatography and size exclusion chromatography in 25 mM MES pH 6.5, 500 mM NaCl and 1 mM DTT.

For endogenous GID^Ant^ purification, yeast strain CRLY45 harbouring 3X Flag tag at the C-terminus of Gid8 was grown in the YPD medium at 30° C and 130 rpm to an OD_600_ of 1. Yeast cells were spun down and rinsed with YP medium to remove the remaining glucose. The pellet was resuspended with YPE medium and grown at 30° C and 130 rpm for 16-19 hours. Next, cell pellets were passed through a 50 ml syringe, to get thin noodle-like pellets and flash-frozen in liquid nitrogen. Frozen yeast noodles were cryomilled using Retsch ZM200 Ultra Centrifugal Mill. Powder was dissolved in the lysis buffer described above. GID complex was purified from the cell lysate by Flag affinity chromatography (Anti-DYKDDDDK G1 resin, GenScript). The resin-bound GID complex was washed with 25 mM MES pH 6.5, 500 mM NaCl, 1 mM DTT, and protein was eluted with 150 μg/ml Flag peptide. The elutions were directly used to make the cryo EM grids.

### Cryo EM sample preparation and Imaging

To prepare cryo EM grids, Vitrobot Mark IV (Thermo Fisher Scientific) was used. 3.5 - 4 μl of freshly purified protein at 0.25 mg/ml was applied to glow discharged Quantifoil holey carbon grids (R1.2/1.3 200 mesh) and incubated for 30 s at 4°C and 100% humidity. Grids were immediately blotted with Whatman no.1 filter paper (blot time 10 s, blot force 10) and vitrified by plunging into liquid ethane.

For GID^SR4^, GID^Ant^, GID^Scaffold^ plus SR^Gid4^, GID^Scaffold^ plus SR^Gid10^, GID^SR4^ minus Gid2/ΔGid9^RING^, GID^SR4^ ΔRINGs and the endogenous GID^Ant^, cryo EM data were collected on a Talos Arctica transmission electron microscope operated at 200 kV and equipped with a Falcon III direct detector. Automated data collection was carried out using EPU software at a nominal magnification of 92,000x, which corresponds to 1.612 Å/pixel at the specimen level, with a total exposure of 63 e^-^/–Å^2^ and the target defocus range between 1.5-3.5 μm.

For GID^SR4^ and GID^Ant^ data were collected on a FEI Titan Krios microscope operated at 300 kV and equipped with a post-column GIF and a K2 Summit direct detector operating in a counting mode. SerialEM software was used to automate data collection (Mastronarde, 2003). Images were recorded at a nominal magnification of 130,000x (1.06 Å/pixel) with target defocus range between 1.1 and 3.2 μm and approximate total exposure of 54 e^-^/Å^2^.

For GID^Scaffold^ plus SR^Gid4^, GID^Scaffold^ plus SR^GID10^, GID^SR4^ minus Gid2/ΔGid9^RING^, images were acquired as described above, except using a K3 direct electron detector instead of K2 and at a nominal magnification of 81,000x corresponding to 1.09 Å**/**pixel at the specimen level. A SerialEM multi-record mode was used to collect data.

### Data processing

Movie frames were motion-corrected and dose-weighted using the Motioncorr2 (Zheng et al., 2017) program. Contrast transfer function parameters were estimated from dose-weighted, aligned micrographs using Gctf (Zhang, 2016). Particles were automatically picked by Gautomatch (https://www.mrc-lmb.cam.ac.uk/kzhang/). Further processing was carried out using Relion (Fernandez-Leiro and Scheres, 2017; Scheres, 2012; Zivanov et al., 2018). Poor quality images were discarded by manual inspection and only particles in the high-quality images were extracted. Iterative rounds of 2D classifications were done to clean up the data. 3D classifications were done using the generated initial model generated and clean set of particles from 2D classification. For large data sets, the particles were split into smaller groups for which 3D classifications were carried out separately, and another round of classification was done if necessary. The resulting 3D classes were manually inspected, and those with complete features were selected for further processing. Particles selected from 3D classification were finally re-extracted, re-centered and subjected to auto-refinement (with and without a mask).

In addition to generating reconstructions for entire complexes, maps with improved quality over specific regions were obtained as follows. A map encompassing the majority of the catalytic module was obtained by multibody refinement of data from GID^Ant^, treating the scaffold module (Gid1-Gid8-Gid5, aka GID^Scaffold^) and the catalytic module (the Gid2-Gid9 subcomplex) as two separate entities. The resultant map over the catalytic module reached 5.1 Å resolution, and enabled visualizing the 4-stranded coiled coil subdomain. Meanwhile, the highest quality map for the CTLH-CRA^N^ domain of Gid9 was obtained using the GID^SR4^ minus Gid2/ΔGid9^RING^ dataset, by focused auto-refinement using a mask over this domain.

Finally, automatic B-factor weighting as well as high resolution noise substitution were done using post-processing in Relion. Local resolution estimates were done as implemented in Relion (Fernandez-Leiro and Scheres, 2017; Scheres, 2012; Zivanov et al., 2018). All the reported resolutions are based on the gold-standard Fourier Shell Correlation (FSC) at 0.143 criterion. Processing details for each dataset are provided in Figure S2, S3, S4A, S4B. Maps generated in this study are summarized in Table S1.

### Model building and refinement

The scheme for model building is shown in Figure S5A, S5B, and described here for each module. The scaffold module, comprising Gid1, Gid8 and Gid5 was built using the 3.4 Å resolution reconstruction of GID^Scaffold^ plus SR^Gid4^. Most of the Gid5, Gid8 as well as SPRY and LisH domains of Gid1 could be built automatically using Buccaneer (Cowtan, 2006), as implemented in CCP-EM software suite (Burnley et al., 2017), with some portions built manually using Coot (Emsley and Cowtan, 2004; Emsley et al., 2010).

The same map was used to build the substrate receptor module, Gid4, manually in Coot (Emsley and Cowtan, 2004; Emsley et al., 2010). The building of Gid4 was guided by a crystal structure of human Gid4 (PDB ID: 6CCR) and sequence alignment of ScGid4 and HsGid4, secondary structure prediction generated by the Phyre^2^ server (Kelley et al., 2015), and the positions of side-chain features (e.g. aromatic residues) as markers.

Segments of Gid9 from the catalytic module were guided by differences in EM reconstructions lacking portions of Gid9. The CTLH-CRA^N^ portion of Gid9 was best visualized and built manually with the 3.5 Å resolution map of Gid9^CTLH-CRA^ generated by focused refinement using data obtained from GID^SR4^ minus Gid2/ΔGid9^RING^. A loop from Gid9 (Gid9^Loop^, residues 291-323) was built using the final 3.2 Å resolution map of GID^SR4^ minus Gid2/ΔGid9^RING^.

Atomic model refinement was performed using ‘phenix.real_space_refine’ available in PHENIX software suite (Adams et al., 2010; Afonine et al., 2018; DiMaio et al., 2013) and the model was validated using Molprobity (Chen et al., 2010). The entire model was checked manually, and regions that lack the sequence registers due to weak/unclear density were modelled as polyalanine. Representative EM density is shown in Figure S5C, and the residues in the final model are summarized in Table S2. Data collection, 3D reconstruction, model refinement and validation details are given in Table 1, 2. Figures of maps and models were prepared with Chimera (Pettersen et al., 2004), ChimeraX (Goddard et al., 2018) and PyMol-v 1.8.2.

### Protein expression and purification for biochemical assays

Insect cell expression as well as cell pellet resuspension for the WT and all the mutant versions of GID^Ant^ and GID^SR4^ used for biochemical assays followed the procedure described in the section **‘**Protein expression and purification for cryo EM’. Proteins were purified from insect cell lysates using Strep-Tactin affinity chromatography by pulling on a Twin-Strep tag fused to Gid8 C-terminus. The eluted proteins were further purified by size exclusion chromatography in 25 mM Hepes pH 7.5, 150 mM NaCl and 1 mM DTT (Buffer B).

To ensure that all assays contained equal concentrations of WT and mutant versions of Gid4 and Gid10 irrespective of their ability to bind GID^Ant^, these proteins that were added exogenously to the *in vitro* assays were expressed as GST-TEV fusions in *E. coli* BL21 (DE3) RIL cells in a Terrific Broth (TB) medium overnight at 20°C. Cell pellets were resuspended in the lysis buffer containing 50 mM Tris-HCl pH 8.0, 200 mM NaCl, 5 mM DTT and 1 mM PMSF (Buffer A). Proteins were purified from bacterial lysates with glutathione affinity chromatography and digested overnight at 4°C with tobacco etch virus (TEV) protease to liberate the GST tag. For further purification, they were subjected to size exclusion chromatography in Buffer B. Remaining free GST as well as uncleaved GST-fusion protein was removed by pass-back over a glutathione affinity resin.

Ubc8, Mdh2 and Mdh2 P2S mutant were expressed in *E. coli* BL21 (DE3) RIL cells in a Terrific Broth (TB) medium overnight at 20°C. Cell pellets were resuspended in Buffer A and proteins were purified from the bacterial lysates by Nickel-Affinity chromatography with Ni-NTA sepharose resin by pulling on the 6xHis tag fused to proteins C-terminus. The elutions were further purified by anion exchange chromatography and size exclusion chromatography in Buffer B.

Untagged WT ubiquitin used for the multi-turnover assays was expressed in *E. coli* BL21 (DE3) RIL cells and purified from bacterial lysates with a glacial acetic acid method (Kaiser et al., 2011). It was further purified by gravity S column ion exchange chromatography and size exclusion chromatography in Buffer B. No-lysine and single-lysine Ub variants as well as WT Ub used for the Ub chain type determination assay were expressed as GST-HRV 3C fusions in *E. coli* BL21 (DE3) RIL cells in a Terrific Broth (TB) medium overnight at 20°C and purified by glutathione affinity chromatography. To liberate the GST tag, elutions were incubated with HRV13 3C protease for 3 hours at room temperature. Further purification was done with size exclusion chromatography in Buffer B that separated Ub from the free GST and uncleaved GST-fusion proteins.

To generate fluorescent Mdh2 (Mdh2-FAM) for ubiquitylation assays, fluorescein was attached to its C-terminus using a sortase A-mediated reaction (Guimaraes et al., 2013). For the reaction, 50 μM Mdh2 fused to a C-terminal sortag (LPETGG) and a 6xHis tag was mixed with 250 μM fluorescent peptide (GGGGG-FAM) and 50 μM sortase A. Reaction was carried out for 30 minutes on ice in a buffer consisting of 50 mM Tris-HCl pH 8.0, 150 mM NaCl and 10 mM CaCl_2_. To get rid of unreacted Mdh2, the reaction mixture was supplemented with 20 mM imidazole and passed through Ni-NTA Sepharose resin. Labelled Mdh2 was purified with size exclusion chromatography in Buffer B.

### Biochemical assays

Unless otherwise stated, *in vitro* ubiquitylation monitored a fluorescently-labelled substrate Mdh2-FAM. All assays were performed at room temperature in a buffer containing 25 mM Hepes pH 7.5, 150 mM NaCl, 5 mM ATP, 10 mM MgCl_2_ and 0.25 mg/mL BSA. At each timepoint, a part of the reaction mixture was quenched by mixing it with SDS-PAGE loading dye. To check the activity of Gid4, Gid5 and Gid1 mutants, as well as to show that Mdh2 ubiquitylation is Gid4 and E2 dependent, the reaction involved mixing of 0.2 μM Uba1, 2 μM Ubc8-6xHis, 1 μM GID^Ant^ (containing either WT or indicated mutants of Gid5 or Gid1), 1 μM Gid4 (Δ1-116; WT or an indicated mutant), 1 μM Mdh2-FAM and 100 μM Ub. For the assay testing importance of the N-terminal meander of Gid4, the exogenously added Gid4 started with the residue at position 80 or 117.

The assay validating dependence of GID^SR4^ activity on the N-terminal proline of its substrate was performed at the same conditions but western blotting with anti-6xHis antibodies was used to visualize ubiquitylation of unlabeled WT and P2S mutant of Mdh2-6xHis (note that complete cleavage of the N-terminal Met residue was confirmed by mass spectrometry).

For testing the mutations in Gid2 and Gid9 RING domains, the assay contained 0.2 μM Uba1, 2 μM Ubc8-6xHis, 1 μM GID^SR4^ (containing either WT or indicated mutants of Gid2 or Gid9), 1 μM Mdh2-FAM and 100 μM Ub. To test the activity of an alternative substrate receptor Gid10, the reaction was composed of 0.2 μM Uba1, 2 μM Ubc8-6xHis, 0 or 1 μM GID^Ant^, 0 or 1 μM Gid10 (either Δ1-56 or Δ1-56 and Δ289-292 version) or 1 μM Gid4 (Δ1-116), 1 μM Mdh2-FAM and 100 μM Ub. Determination of the type of Ub chain formed by GID^SR4^ was done by using a panel of single-Lys Ub variants, with all other lysines mutated to arginines. The reaction mixture was composed of 0.2 μM Uba1, 2 μM Ubc8-6xHis, 1 μM GID^SR4^, 1 μM Mdh2-FAM and 20 μM Ub (WT, lysineless K0 Ub or any of the single Lys Ub variants).

To analyze if addition of the substrate receptor Gid4 to the GID^Ant^ has any impact on its intrinsic E3 ligase activity, a substrate-independent discharge assay was employed. To separate an effect of E2∼Ub discharge from its E1-dependent loading, this assay was performed in a pulse-chase format. In the pulse reaction, loading of E2 was performed by mixing 0.5 μM Uba1, 10 μM Ubc8-6xHis and 30 μM Ub in a buffer containing 25 mM Hepes pH 7.5, 100 mM NaCl, 1 mM ATP and 2.5 mM MgCl_2_. After 15 minutes of incubation of the pulse mixture at room temperature, E2 loading was stopped by addition of 50 mM EDTA. For the chase reaction, the quenched pulse mixture was mixed with an equal volume of the chase initiating mixture containing 1 μM GID^Ant^, 0, 1 or 5 μM Gid4 (Δ1-116) and 40 μM lysine pH 8.0 in 25 mM Hepes pH 7.5 and 100 mM NaCl, and incubated at room temperature. The discharge was quenched at each of the timepoints by mixing the discharge reaction with SDS-PAGE loading dye without any reducing agent and visualized with a non-reduced SDS-PAGE stained with Coomassie.

To test if our recombinant Mdh2 binds to Gid4 according to the Pro/N-degron pathway, purified GST-tagged Gid4 (Δ1-116) was mixed with two-fold molar excess of Mdh2-6xHis (WT or the P2S mutant) in a buffer containing 25 mM Hepes pH 7.5, 150 mM NaCl and 1 mM DTT. After incubating the proteins for 30 minutes on ice, 20 μL of GST resin was added to the mixture and further incubated for 1 hour. As a negative control, Mdh2-6xHis was mixed with GST resin in absence of Gid4. GST beads were then thoroughly washed and proteins were eluted. The elution fractions were analyzed with SDS-PAGE to check for the presence or absence of an Mdh2 band. A similar binding test was applied to check if an alternative substrate receptor Gid10 interacts with GID^Ant^. Here, GID^Ant^, which was Twin Strep-tagged on Gid8 C-terminus, was mixed with a two-fold molar excess of Gid10 (Δ1-56) and Strep Tactin pull-down was performed. As a negative control, Gid10 was mixed with Strep Tactin resin in the absence of GID^Ant^.

## QUANTIFICATION AND STATISTICAL ANALYSIS

For the *in vivo* Fbp1 degradation assay, experiments were performed in at least biological triplicate. Fbp1 degradation pattern was visualized by western-blot and the bands were quantified. Bars on graphs represent average (n>=3) and error bars represent standard deviation.

